# A two-step organoid screen identifies transient p53 activation as a senolytic strategy for radiation-induced salivary gland dysfunction

**DOI:** 10.64898/2026.09.29.754812

**Authors:** Abel Soto-Gamez, Yi Wu, Eline Hageman, Uilke Brouwer, Davide Cinat, Arjan Vissink, Marco Demaria, Lara Barazzuol, Sarah Pringle, Rob P. Coppes

## Abstract

Radiotherapy significantly improves the survival of head and neck cancer patients. However, it frequently damages the salivary glands, causing persistent hyposalivation and xerostomia. Radiation-induced cellular senescence within the salivary gland stem/progenitor cell niche contributes to hyposalivation, suggesting that selective elimination of senescent cells could promote tissue recovery. However, the effectiveness of senolytic compounds is strongly context dependent, varying with tissue of origin and potentially impairing regenerative cells required for functional restoration. Here, we developed a two-step organoid-based screening methodology that integrates the preferential induction of apoptosis in irradiated, senescence enriched, salivary gland organoids with the preservation of stem/progenitor cell regenerative capacity. As a first step, the senolytic activity of candidate drugs was tested to identify drugs that induced apoptosis in senescent cells of irradiated organoids but were not or less toxic to unirradiated organoids. As a second step, the confirmed senolytic compounds were tested for effects on stem cell self-renewal potential. Using this approach, we identified Nutlin3a, an MDM2 inhibitor and p53 activator, as the compound with the most favorable toxicity profile in salivary glands. Last, we confirmed the anti-senescence and pro-regenerative potential of Nutlin3a in an *in vivo* mouse model of radiation-induced hyposalivation, where localized Nutlin3a delivery resulted in a robust increase in saliva production, as early as two weeks after administration. These findings establish an organoid-based strategy for identifying tissue-adapted senolytic interventions and propose transient local p53 activation with Nutlin3a as a potential approach to mitigate radiation-induced salivary gland dysfunction.

## INTRODUCTION

Radiotherapy is the standard of care for many head and cancers, significantly improving patient survival. However, the irradiation of normal tissue surrounding the tumor is unavoidable and often includes the salivary glands [1]. Chronic hyposalivation and consequential xerostomia are common and debilitating side effects of salivary gland irradiation, leading to numerous complications that severely affect patient quality of life. These include impaired speech, taste, difficulty swallowing, ulceration of the oral mucosa, and severe dental complications requiring extraction [2, 3]. Despite improvements in radiotherapy delivery, there are no effective interventions to restore salivary gland function after radiation injury.

Cellular senescence has emerged as one mechanism contributing to persistent radiation-induced tissue dysfunction. Senescent cells undergo a durable proliferative arrest and acquire broad phenotypic changes, including altered survival signaling and a senescence-associated secretory phenotype (SASP) that can modify neighboring cells and the tissue microenvironment [6]. We and others have previously shown that irradiation induces senescence within the salivary gland stem/progenitor cell niche, and that reducing the burden of senescent cells using genetic and pharmacological models improves regenerative capacity and salivary gland function [4, 5]. These observations provide a rationale for senolytic therapy, compounds that can preferentially induce senescent cell death, after radiotherapy.

A major challenge, however, is that senolytic vulnerabilities are not universal. Senescent cells remodel multiple pro- and anti-apoptotic pathways, and the survival dependencies that emerge vary according to cell type and senescence-inducing stress [6]. Consequently, compounds that are senolytic in one experimental context may be ineffective or toxic in another. For example, adaptive responses can render senescent cells more resistant to challenges with certain pro-apoptotic chemotherapeutics compared to non-senescent ones, such as staurosporine [7], thapsigargin [8], FasL [9, 10], TRAIL [11], and many others. However, the reliance of senescent cells on anti-apoptotic pathways for cell survival can conversely render them more susceptible to pharmacological approaches targeting these pathways. For regenerative tissues, selectivity is particularly important as preferential killing of damaged or senescent cells is unlikely to provide durable benefit if the same treatment compromises the stem/progenitor cells required for tissue repair. The identification of novel effective and less or non-toxic senolytic drugs is, therefore, an unmet need that could benefit from the development of novel screening methods.

Organoid models provide an opportunity to address both requirements in a tissue-relevant system. Salivary gland organoids retain stem/progenitor cell activity and can be used both to quantify treatment-induced cell death and to measure subsequent organoid-forming capacity as a functional readout of regeneration [12, 13]. We therefore reasoned that candidate senolytics should be evaluated sequentially for preferential induction of apoptosis in irradiated, senescence-enriched organoids, but also for preservation or recovery of stem/progenitor cell function.

Here, we establish this two-step screening strategy in a model of radiation-induced salivary gland injury. The screen identified the MDM2 inhibitor Nutlin3a as a candidate with preferential toxicity toward irradiated organoids and a favorable regenerative profile. We then investigated the apoptotic state associated with this selectivity and tested whether local Nutlin3a administration could improve salivary gland function after irradiation in vivo.

## MATERIALS AND METHODS

### Mice

C57BL/6 mice (Envigo, Harlan, The Netherlands) were housed in the central animal facility of the University Medical Center Groningen under conventional conditions. All animal procedures were approved by the Central Committee of Animal Experimentation of the Dutch government and by the Institute Animal Welfare Body of the University Medical Center Groningen.

### Isolation of murine salivary gland cells and organoid culture

Submandibular salivary glands (SGs) were dissected from 8-to 12-week-old female mice, mechanically and enzymatically dissociated to single cells, and cultured in DMEM:F12 medium (Gibco, The Netherlands) containing 1% penicillin / streptomycin (Gibco) and 2 mM GlutaMAX (Thermo Fisher Scientific, The Netherlands). The medium was supplemented with EGF (20 ng/mL, Sigma-Aldrich, The Netherlands), FGF-2 (20 ng/mL, PeproTech, NJ, USA), insulin (10 µg/mL, Sigma-Aldrich), dexamethasone (1 µM, Sigma-Aldrich), and N2 supplement (1× Gibco). After 3 days in culture, primary salispheres were dissociated to single cells and embedded in Matrigel domes. After solidification, the domes were covered with the medium described above supplemented with the ROCK inhibitor Y-27632 (10 µM, Abcam, UK), 10% R-spondin1 conditioned medium, and 50% Wnt3a conditioned medium, referred to as WRY medium. For routine culture SG organoids were passaged weekly by dissociation and reseeding in WRY medium.

### SG organoid imaging-based screening

Single cells derived from SG organoids were seeded in multi-well plates. For 96 well plates, 2500 cells/ well were embedded in 20 µl of Matrigel:WRY medium (2:1). After gel solidification, 200 µL of WRY medium was added. After 5 days in culture, organoids were next sham-irradiated or irradiated, the medium was refreshed, and cultures were maintained for an additional 7 days to allow the radiation-induced phenotype to develop. Organoids were then treated with the indicated drugs and concentrations (Table 1) in WRY medium containing 1 µM CellEvent Caspase-3/7 Green Detection Reagent (Thermo Fisher Scientific). Plates were immediately transferred to an IncuCyte S3 (Essen BioScience) and maintained at 37 °C and 5% CO2. Green fluorescence was recorded by IncuCyte® ZOOM every 2 hours under 4 × magnification for up to 48 hours. The area under the curve (AUC) was calculated for each drug under both conditions (non-irradiated vs. irradiated) in GraphPad Prism 8.0 (Baseline Y=0, peak definition above baseline). The senolytic index was defined as AUC_IR_ – AUC_Non-IR_.

### RNA isolation and RT-qPCR

Total cellular RNA was isolated using a commercial kit according to the manufacturer’s instructions (Qiagen, RNeasy Mini Kit, Hilden, Germany). The concentrations of mRNA were measured by NanoDrop ND-100 spectrophotometer (Thermo Fisher Scientific, Waltham, MA, USA). cDNA was obtained through a reverse transcription reaction using a Reverse Transcription Kit (Promega, Madison, WI, USA). A panel of senescence-associated genes consisting of mouse Cdkn2a/p16, Tnfrsf10, Gdnf and Cxcl10, p53 target genes, and pro- and anti-apoptotic genes, were detected by RT-qPCR using a commercially available SYBR assay kit (Bio-Rad). mRNA levels of the YWHAZ gene were used as a reference for data normalization. The sequences of primers were listed in Table 2.

#### Bulk RNA-sequencing analysis

Bulk RNA-seq data from (sham)-irradiated SG organoids was analyzed from a previously described dataset [14]. Briefly, fastq files were aligned with the mouse genome GRCm39, and R (v4.3.2) and RStudio were used for downstream analyses. The R package edgeR was used for normalization and to identify differentially expressed genes (LogFC > 0.6, FDR < 0.05). Lowly expressed genes were excluded from the analysis (total count <1 in less than 2 samples).

### Western Blotting

Organoids were harvested after drug treatment and homogenized by 2 times 5 second sonication in Laemnli buffer. Protein concentration of the homogenates was determined by Bradford quantification method. Cell lysates were dissolved by 2x SDS sample loading buffer (20% SDS, 50% glycerol, 0.33 M Tris-Hcl pH = 6.8 and 0.05% bromophenol blue) and 10% β-mercaptoethanol and then boiled at 99 ℃ for 5 min. Equal protein amount was separated with 12.5% SDS-PAGE gels (BIO-RAD) and transferred to PVDF membranes using Trans Blot Turbo System (BIO-RAD). The membranes were blocked in 10% milk in PBST and incubated for 1 h. For immunoblots, the primary antibodies were used: mouse anti-Bcl-2 (1:1500, sc-7382, Santa Cruz Biotechnology), rabbit anti-Survivin (1:2000, 10508-1-AP, Proteintech) and mouse anti-GAPDH (1:10000, Fitzgerald). Blots were subsequently incubated with HRP-conjugated anti-rabbit (1:5000, Amersham, NA934V) or anti-mouse (1:5000, NXA931V, Amersham) secondary antibodies and the membranes were developed using ECL reagent (Thermo Fisher Scientific). The protein signals were analyzed by Image Lab software.

### SA-β-galactosidase (gal) staining in organoids

Organoids were collected after drug treatment, fixed, and whole-mount stained for 4 h with SA-β-gal detection solution (pH = 6.0) according to the manufacturer’s instructions (Merck Millipore, KAA002, MA, USA). Positive cells were identified as blue-stained cells under light microscopy.

### In vivo irradiation and localized treatment with Nutlin3a

The experimental setup for *in vivo* irradiation was designed as previously described [4]. In short, the salivary glands of 2–3-month-old female C57BL/6 mice were (sham-)irradiated with a single dose of 15 Gy X-rays (Precision X-ray). Eight weeks after irradiation, mice were treated with vehicle (high purity corn oil, Cat. No. C8267, Sigma Aldrich) or with Nutlin3a (0.05 mg/kg) (Cat. No HY-10029, MedChemExpress) dissolved in vehicle and delivered locally by retroductal instillation/infusion to each gland in atropine (0.5 mg/kg) treated mice as previously described [15], followed by a subsequent dose delivered to each gland after a 2-week rest period. Saliva production was measured as previously described [16]. Briefly, saliva was collected for 15 min after stimulation with pilocarpine (2 mg/kg). Measurements were made at baseline (1 and 2 weeks prior to irradiation), and 7, 13, and 22 weeks after irradiation and/or Nutlin3a treatment. Mouse were sacrificed at 13 weeks or 22 weeks after irradiation. Sham-irradiated controls (n=3) were complemented with historic data obtained from Peng et al. [4] to reduce the number of animals sacrificed where no effect was expected

### Immunohistochemistry and immunofluorescent staining

Salivary glands were fixed immediately after extraction with 4% formaldehyde for 24 hours and embedded into paraffin. Five micrometer paraffin sections were deparaffinized and boiled for 8 min with pre-heated antigen retrieval buffer. Next, sections were incubated with primary antibodies for rabbit anti-P21 antibody (sc-471, Santa Cruz Biotechnology, TX USA), or rabbit anti-Aquaporin5 antibody (Alomone labs, AQP-005, Jerusalem, Israel), or mouse anti-GLB1 β-gal antibody (4F4F4, Thermo Fisher Scientific, The Netherlands). Visualization for bright field microscopy was accomplished by adding specific secondary biotin carrying antibodies, biotinylated rabbit anti-mouse (Dako, E0413), or biotinylated Swine anti-rabbit (Dako, E0431) at 1:300 dilution. Nuclear counterstaining was performed with hematoxylin. The percentage of area positive for AQP5 staining in salivary gland tissue was quantified using ImageJ on five representative fields at 10X magnification on at least three mice per group. For mouse anti-γH2AX (Merck, 06-636, Darmstadt, Germany).

### Statistical analysis

GraphPad Software (v8.0.1) was used for statistical analyses. Values are presented as mean ± SEM from at least 3 independent experiments, unless stated otherwise. Group sizes are provided in the figure legends. Two-tailed *t*-tests or two-way ANOVA were used as indicated. *P* values < 0.05 were considered statistically significant.

## RESULTS

### Radiation remodels apoptotic signaling and creates a first-step platform for tissue-specific senolytic screening

As a starting point, we leveraged the use of organoid models to verify the selectivity of senolytic drugs in a specific tissue. Using the salivary gland as an example, we adapted our previously developed three-dimensional organoid model consisting of *ex vivo* expanded salivary gland stem/progenitor (SGSC) cells [12]. When grown using 3D culture methods, single SGSCs are capable of forming organoids/miniglands containing all salivary gland cell lineages and are a useful *in vitro* model to study salivary gland regeneration [13, 17].

We first asked whether irradiation produces an apoptotic state that could reveal therapeutically exploitable survival dependencies in salivary gland cells. Probing single cell transcriptomics datasets of mouse salivary gland tissue [18] and organoids [14], we observed an increased expression of senescence-associated markers, including Cdkn1a, Cxcl1, and Tnfrsf10 after irradiation (**Fig. 1A**), together changes in apoptosis-related signaling (**Fig. S1A,B**). Indeed, senescent cells employ various genes in the mediation of apoptosis resistance, in a cell-type dependent manner. Bulk RNA sequencing of non-irradiated and irradiated (7 Gy) SG organoids further revealed the simultaneous induction of pro- and anti-apoptotic genes (**Fig. 1B**). Among the induced pro-apoptotic genes were the BH3-only factors Bmf and Bik [19, 20]. Several P53 pathway-related genes, well-established to be associated with apoptosis, were also identified including Trp53inp2, Trp53cor1, Cdkn1a, Bbc3, Mdm2 (a negative regulator of P53), and Tnfsf10/TRAIL (an extracellular ligand that acts as a potent inducer of extrinsic apoptosis). Conversely, also anti-apoptotic genes were upregulated, including Tnfrsf11b (which functions as a soluble decoy receptor for TRAIL), and the Bcl-2 family members Mcl-1 and Bcl2l1. Gene-set enrichment analysis independently supported enhanced apoptotic signaling after irradiation (**Fig. S1C**). Thus, radiation did not simply suppress apoptosis but also established a remodeled state containing concurrent death-promoting and survival signals

**Figure 1.**
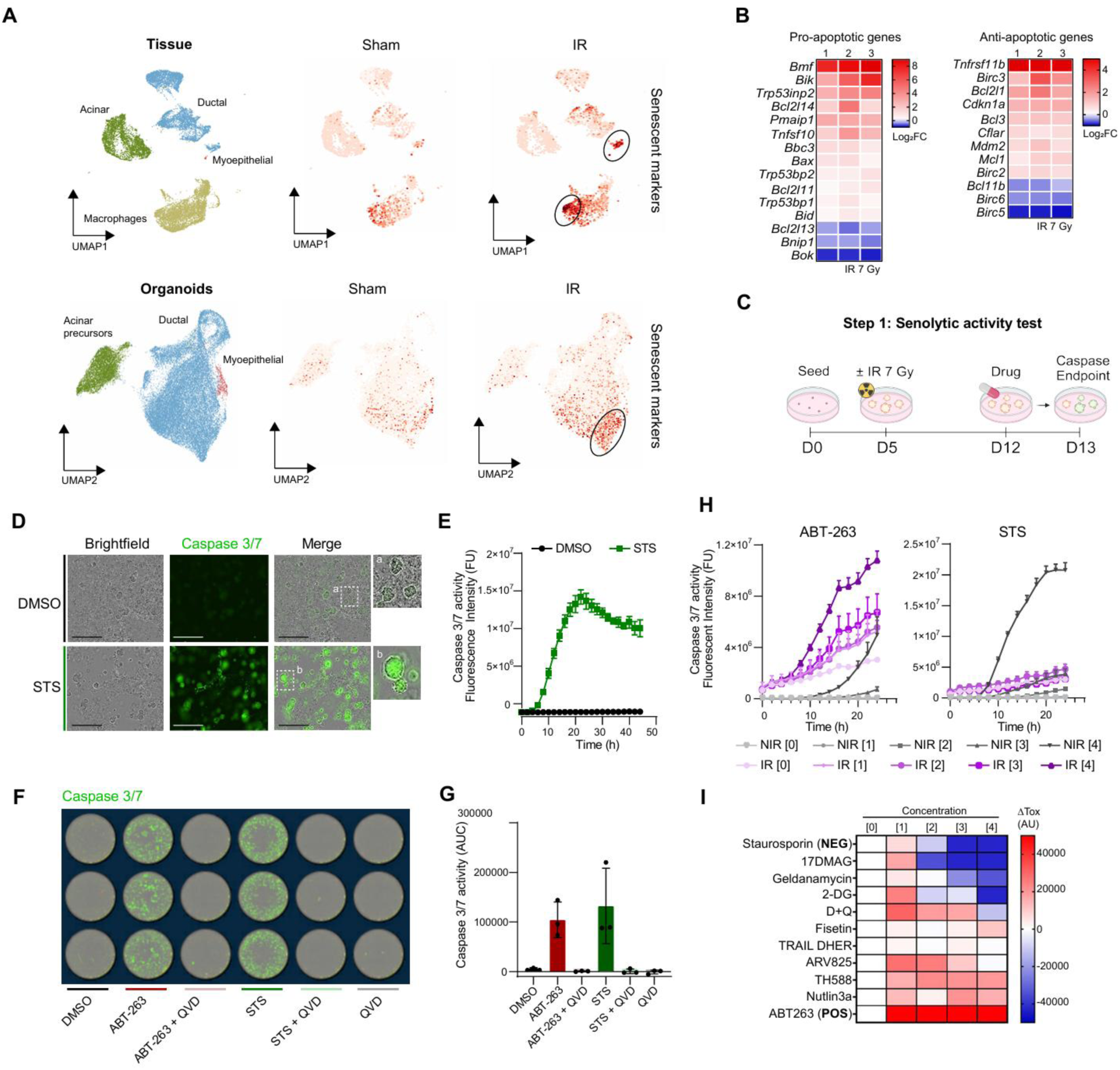
Radiation remodels apoptotic signaling and enables a first-step organoid-based screen for preferential cell killing. (**A**) Expression of senescence-associated markers (Cdkn1a, Cxcl1, Tnfrsf10) in single-cell transcriptomic datasets from sham-irradiated and irradiated salivary gland tissue [18] and organoids [14] (10 Gy and 7 Gy, respectively). (**B**) Differential expression of pro- and anti-apoptotic genes of irradiated SG organoids relative to non-irradiated organoids by bulk RNA sequencing. (**C**) Schematic of the caspase-3/7 fluorogenic assay. (**D, E**) Fluorescence response of organoids treated with staurosporine (STS, 10 µM) as a positive control for apoptosis induction (Caspase-3/7 positive cells are shown in green, scale bar: 100 µm). (**F-H**) Validation of apoptosis-dependent fluorescence using staurosporine and ABT-263, with QVD-OPh as a pan-caspase inhibitor. (**H**) Time-resolved comparison of caspase activity in irradiated (IR) and non-irradiated organoids (NIR) treated with ABT-263 or Staurosporin confirming differential responses. (**I**) Senolytic index calculated as AUC_IR_ – AUC_NIR,_ across candidate compounds and concentrations. A positive value (red color) indicates higher toxicity in irradiated (senescent) samples (e.g. ABT-263, positive control “POS”), whereas a negative value (blue color) indicates a higher toxicity in non-irradiated (non-senescent) samples (e.g. Staurosporine, negative control “NEG”).

To investigate the toxicity of potential senolytics drugs on healthy, non-irradiated SG cells, we first focused on the visualization of apoptosis using a caspase-cleavable fluorogenic probe. Caspases are synthesized as inactive zymogens and their activation is a tightly regulated process. In particular, caspase cascades are commonly initiated by caspases 8 and 9, and converge in the activation of caspases 3 and 7 [21], responsible for the direct cleavage of key cellular proteins in the degradative stage of apoptosis. Consequently, the Caspase 3/7 peptide substrate Asp-Glu-Val-Asp (amino acid abbreviation: DEVD) has been used extensively in caspase-cleavable probes for imaging apoptosis [22]. Here, we employed a DEVD peptide sequence conjugated to a quenched nucleic acid-binding dye. In the presence of active caspase 3/7, the peptide is cleaved, releasing the dye which can then bind to nuclear DNA and produce a fluorogenic response indicative of apoptosis (**Fig. 1C**). The caspase-3/7 fluorogenic substrate was tested with staurosporine as a positive control (**Fig. 1D**), a broad-spectrum kinase inhibitor typically used for the induction of apoptosis upstream of DEVD-specific caspase activation [23]. As expected, a time-dependent increase in fluorescence intensity was observed upon treatment of 5-day-old SG organoids with staurosporine (1 µM) with a maximum signal observed around 24 hours post-treatment, after which a plateau was reached (**Fig. 1E**). The dependence of the fluorogenic response on active caspases was demonstrated by the co-treatment with the pan-caspase inhibitor QVD-Oph (QVD, 10 µM), where no increase in fluorescence signal was observed (**Fig. S2A**).

The selectivity of the DEVD probe was confirmed in a 96 well format using a whole well scanning technique to measure differences in fluorescence signals in organoids treated with ABT-263 (a Bcl-2 inhibitor that induces apoptosis) or staurosporine (**Fig. 1D,E**). Expectedly, increased fluorescence was observed when organoids were treated with either drug, but the signal was completely abolished when organoids were treated in the presence of QVD-Oph (**Fig. 1F, G**). Similar findings were observed when propidium iodide incorporation was used as an alternative readout for cell death (**Fig. S2B**).

To verify preferential toxicity of candidate senolytic agents against radiation-induced senescent cells, as opposed to non-irradiated cells, we measured caspase activity in sham- and irradiated SG organoids, with and without senolytic agent treatment. For this, five-day-old (D5) organoids were irradiated with 7 Gy and analyzed 7 days later (D12), a dose and time known to induce senescence in SG organoids [4].

We have previously shown an effective but modest improvement of saliva production post-irradiation after systemic treatment with the senolytic drug ABT263 (a Bcl-2 family inhibitor) [4]. Therefore, (sham-)irradiated SG organoids were treated with ABT-263 as a positive control for induction of cell death in salivary gland senescent cells. Changes in fluorescence were tracked every 2 hours for a total period of 24 hours. As shown in **Fig. 1H**, increased toxicity was observed in irradiated cells upon treatment with ABT-263, as shown by a higher increase in caspase activity, which was also evident at earlier timepoints compared to non-irradiated organoids (**Fig. 1H)**. In contrast, staurosporine was less effective at inducing cell death in irradiated organoids, in line with reports of resistance to staurosporine-induced apoptosis in senescent cells [7]. We expanded the panel to include additional drugs previously described as senolytics and evaluated their effectiveness as senolytics in the salivary gland [4, 11, 24–28]. The panel of drugs was tested at 4 different concentrations (**Supplementary Table 1, Fig. S3**) and a ‘senolytic index’ describing toxicity in irradiated samples relative to unirradiated was calculated. Drugs were considered to have senolytic properties in SG organoids when they preferentially induced apoptosis in irradiated organoids (‘senolytic index’ higher than 0, shown in color red, **Fig. 1I**).

As observed in **Figure 1I**, the positive control ABT-263 had a positive senolytic index, whereas the negative control staurosporine had a negative senolytic index. Several of the drugs tested induced higher caspase activity in irradiated organoids compared to non-irradiated organoids (i.e. had a positive senolytic index: D+Q, TH588, ARV825, Nutlin3a), suggesting senolytic activity in salivary gland cell types, whereas other drugs were more toxic to non-irradiated organoids, suggesting a lack of senolytic activity in SG organoids (i.e. negative senolytic index: 17DMAG, Geldanamycin, 2DG).

### A functional second-step screen prioritizes Nutlin3a by coupling selective cell death to regenerative capacity

Preferential killing of irradiated cells is not sufficient for a regenerative therapy if the treatment also damages the surviving stem/progenitor compartment. We therefore introduced a second screening step in which compounds with a positive senolytic index were evaluated for their effect on organoid-forming efficiency (OFE), a functional measure of salivary gland stem/progenitor cell activity. Organoids were irradiated on day 5 and maintained until day 12 to allow the radiation-induced phenotype to develop, in agreement with our previous studies [4]. We selected doses of ABT-263, D+Q, TH588, ARV825, and Nutlin3a that showed preferential activity in the first screen (**Supplementary Table 1**). After 24 hours of treatment, organoids were dissociated to single cells and replated to quantify OFE (**Fig. 2A**).

**Figure 2.**
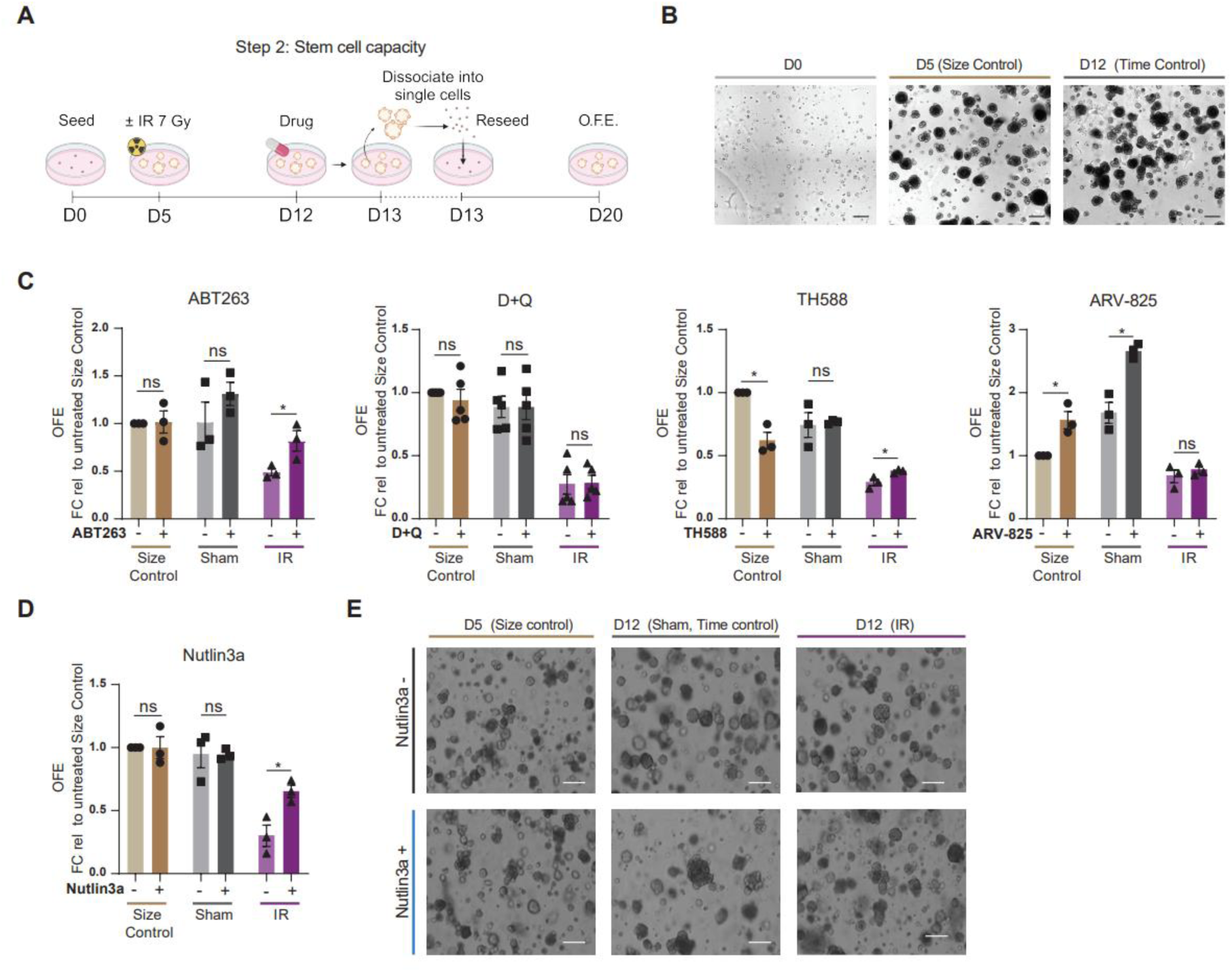
A functional second screen identifies Nutlin3a as the lead candidate to restore the regenerative potential of irradiated salivary glands. (**A**) Experimental design for measuring organoid forming efficiency (OFE) after drug candidate treatment. (**B**) Irradiated organoids were treated with candidates on day 12, dissociated after 24 hours, and replated as single cells, Day-5 sham organoids were used as a size-matched control, and Day-12 sham organoids were used as a time-in-culture control. (**C**) Organoid forming efficiency (OFE) was calculated after treatment with candidate compounds at doses selected from the apoptosis screen. (**D**) Nutlin3a produced the largest recovery of OFE in irradiated organoids, while showing a favorable profile in non-irradiated controls. (**E**) Representative images of organoids after Nutlin3a treatment are shown (Scale bar, 200 μm). Data shown as mean ± sem from *N* = 3/group, ns = not significant, \**p* < 0.05, in unpaired *t*-test.

Because irradiation strongly restricts organoid growth, two sham-irradiated controls were included. Day-5 sham organoids served as a size-matched control for day-12 irradiated organoids, whereas day-12 sham organoids controlled for time in culture and the effects of prolonged culture stress (Fig. 2B). This design allowed us to distinguish treatment effects on irradiated cultures from nonspecific effects on proliferating, non-irradiated cells.

ABT-263, TH588, and Nutlin3a increased OFE after irradiation by approximately 1.7-, 1.3-, and 2.2-fold, respectively (**Fig. 2C-E**). Nutlin3a produced the largest recovery and showed the most favorable profile across irradiated and control conditions. By contrast, TH588 also reduced OFE in the size-matched non-irradiated control, indicating toxicity toward the self-renewal potential of healthy proliferative cells. We therefore selected Nutlin3a for further validation.

### Nutlin3a preferentially induces apoptosis in irradiated, senescence-enriched salivary gland cells

Based on the findings of the two screening steps, we decided to focus on validating the senolytic effect of Nutlin3a. For this, we tested the effect of Nutlin3a on (sham)-irradiated SG organoids at a broader range of concentrations, where we confirmed optimum senolytic activity when tested at concentrations around 10 µM in both caspase activity (**Fig. 3A**) and cell viability in an MTS assay (**Fig. 3B**). Accordingly, we observed a leftward shift in the IC50 curve of Nutlin3a in irradiated organoids compared to sham, thereby confirming a higher toxicity of Nutlin3a after irradiation (**Fig. 3C**).

**Figure 3.**
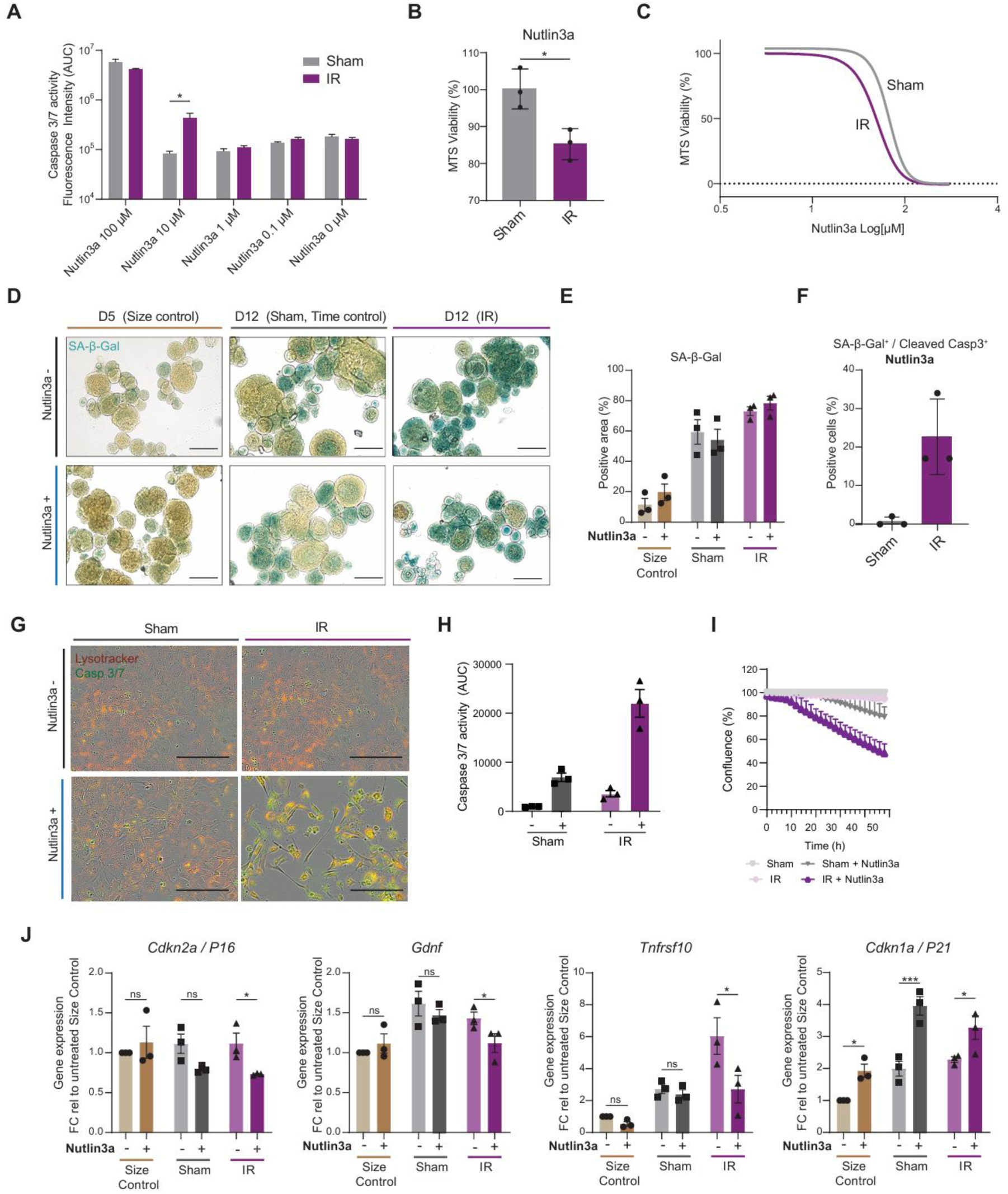
Nutlin3a preferentially induces apoptosis in irradiated, senescence-enriched salivary gland cells. (**A**) Caspase-3/7 activity and (**B**) cell viability after Nutlin3a treatment across a concentration range in sham-irradiated and irradiated SG organoids. (**C**) Dose-response analysis showing increased sensitivity of irradiated cells. (**D**) SG organoids were fixed and stained for SA-β-gal (blue) after treatment with Nutlin3a, representative images of are shown. Scale bar, 100 μm. (**D,E**) SA-β-gal staining after Nutlin3a treatment. (**F**) Cleaved Caspase-3 within SA-β-gal positive cells. (**G-H**) Preferential toxicity of Nutlin3a in irradiated SG cell monolayers after 48 hours (**J**) Expression of senescent-associated marker gene expression in SG organoids after Nutlin3a treatment. Data shown is mean + sem from *N* = 3/group, ns = not significant, \**p* < 0.05 in an unpaired *t*-test.

Next, we assessed the effect of Nutlin3a treatment on known senescence markers. First, we measured senescence-associated beta-galactosidase (SA-β-gal) in (sham-)irradiated organoids. However, we failed to see a clear change in SA-β-gal staining after Nutlin3a treatment (**Fig. 3D, E**), despite increased cleaved caspase 3 in SA-β-gal cells following Nutlin3a treatment in irradiated samples (**Fig. 3F**), likely due to their physical presence in the organoid after apoptosis. The selectivity of Nutlin3a was next confirmed in 2D cultured (sham)-irradiated salivary gland cells (**Fig. 3G**), where Nutlin3a treatment led to higher increases in caspase activity (**Fig. 3H**), and a lower confluence (**Fig. 3G**) after drug treatment for 48 hours.

To further investigate the senolytic properties of Nutlin3a in salivary gland cell types, we measured the senescence-associated cell cycle arrest markers Cdkn1a/P21 and Cdkn2a/P16, the membrane marker Tnfrsf10, and the SASP protein Gdnf in (sham)-irradiated organoids (**Fig. 3I**). Of these, a decrease in all senescence markers was observed in irradiated organoids treated with Nutlin3a when compared to untreated, except for the Cdkn1a/P21, possibly due to its role as a downstream target of p53. These findings suggest Nutlin3a decreases but not totally ablates the senescent cell burden in irradiated salivary glands.

### Local Nutlin3a treatment produces functional recovery after salivary gland irradiation

Nutlin3a was first identified as a potent and selective small-molecule antagonist of MDM2 in its p53-binding pocket [29], thereby inhibiting the interaction between the two proteins (MDM2 and p53), and resulting in the activation of the p53 pathway. As expected, we observed a transcriptional upregulation of p53 target genes such as P53INP1, MDM2, after Nutlin3a treatment (**Fig. 4A**).

**Figure 4.**
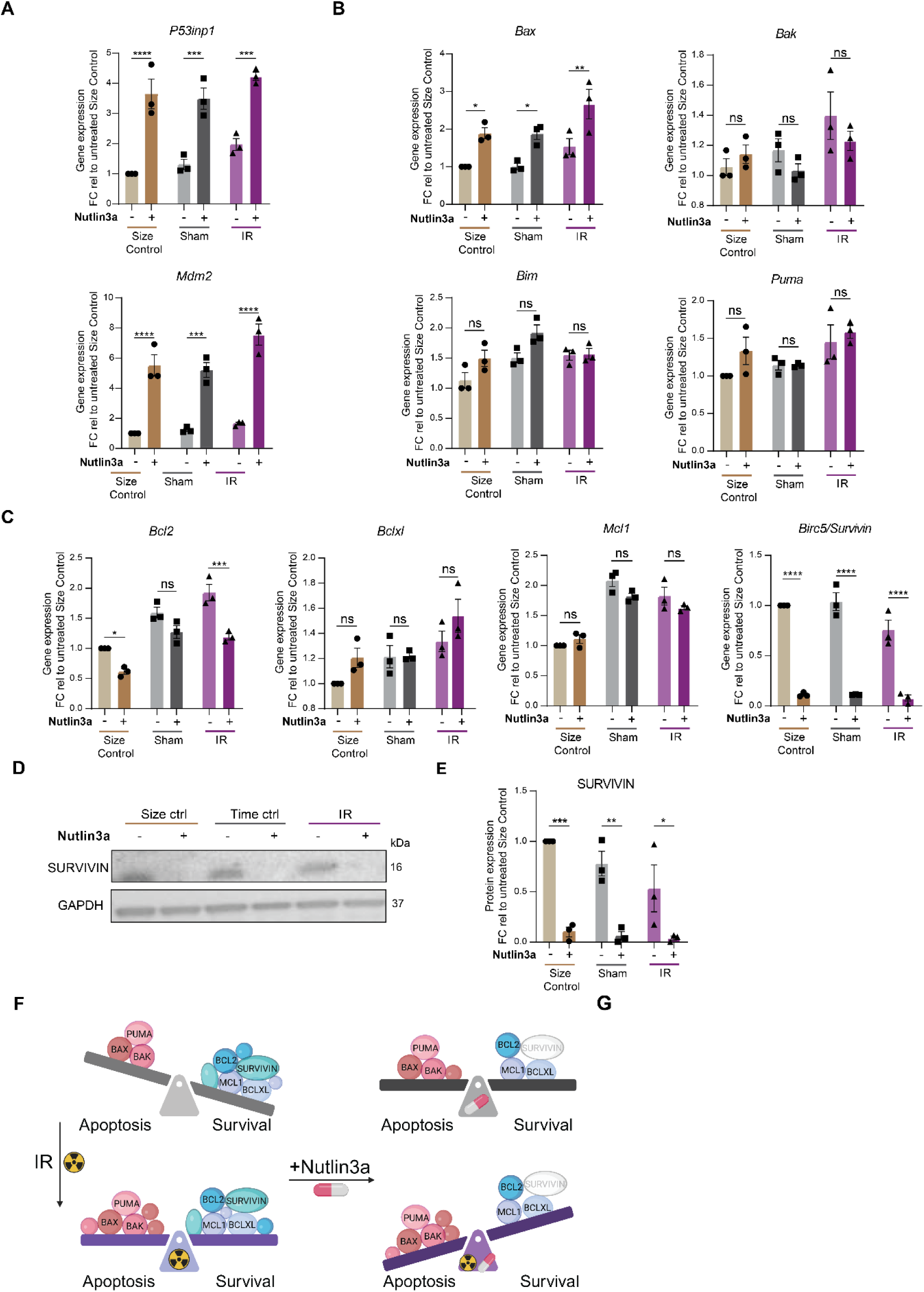
Nutlin3a shifts apoptosis-primed irradiated salivary gland cells toward cell death. SG organoids were treated with Nutlin3a for 24 hours and analyzed for (**A**) p53 target genes, (**B**), pro-apoptotic genes, (**C**) and anti-apoptotic genes by RT-qPCR. (**D**) Reduction of the anti-apoptotic gene Birc5/Survivin after Nutlin3a treatment was confirmed at the protein level by immunoblotting and (E) band intensity-quantification normalized to the untreated size control (n = 3/group). (**F**) Proposed model in which irradiation establishes an apoptosis-primed state and Nutlin3a shifts the balance toward cell death through p53-dependent changes in pro- and anti-apoptotic signaling. Data shown are mean ± sem, ns = not significant, \**p* < 0.05, \*\**p* < 0.01, \*\*\**p* < 0.001, \*\*\*\**p* < 0.0001, unpaired *t*-tests.

Interestingly, the upregulation of p53 target genes following Nutlin3a treatment was observed not only in irradiated organoids, but also in non-irradiated organoids collected after 5 days in culture (size controls), and in 12 days in culture (time controls, sham irradiated). However, given the preferential toxicity observed in irradiated organoids (**Fig 4A-B**), we next assessed the transcriptional levels of both pro-apoptotic and anti-apoptotic genes across conditions.

Among the pro-apoptotic genes analysed, Bax showed the strongest and most consistent transcriptional upregulation upon treatment with Nutlin3a, with mRNA levels significantly elevated across conditions (size control D5, time control D12, and irradiated organoids IR-D12) (**Fig. 4B**). In contrast, a transcriptional downregulation of the anti-apoptotic genes Bcl2 and Birc5 gene, which codes for the protein Survivin was observed after Nutlin3a treatment (**Fig. 4C**). Of the anti-apoptotic genes examined, Birc5/Survivin was consistently downregulated across all conditions tested following Nutlin3a. A significant decrease of Birc5/Survivin was also confirmed at the protein level using Western Blotting (**Fig. 4D-E**). Of interest, a protective, anti-apoptotic role of Survivin has been previously proposed in aged fibroblasts [30], and (doxorubicin) therapy-induced senescent cells [31]. Altogether these findings and other literature suggest that the higher levels of pro-apoptotic proteins observed post-irradiation (such as Bax), paired with a downregulation of anti-apoptotic proteins after Nutlin3a treatment (such as Survivin), may result in the increased cell death of senescent salivary gland cells, but reduced toxicity to non-irradiated cells (**Fig. 4F**).

### Local Nutlin3a treatment produces functional recovery after salivary gland irradiation

Next, we decided to test *in vivo* whether Nutlin3a treatment improved SG function after irradiation. For this we adapted our previously described radiation-induced hyposalivation model where C57BL/6 mice are locally irradiated with a single dose 15 Gy delivered to the salivary gland neck region, followed by senolytic intervention at 8 and 10 weeks post-irradiation (**Fig. 5A**), as previously described [4].

**Figure 5.**
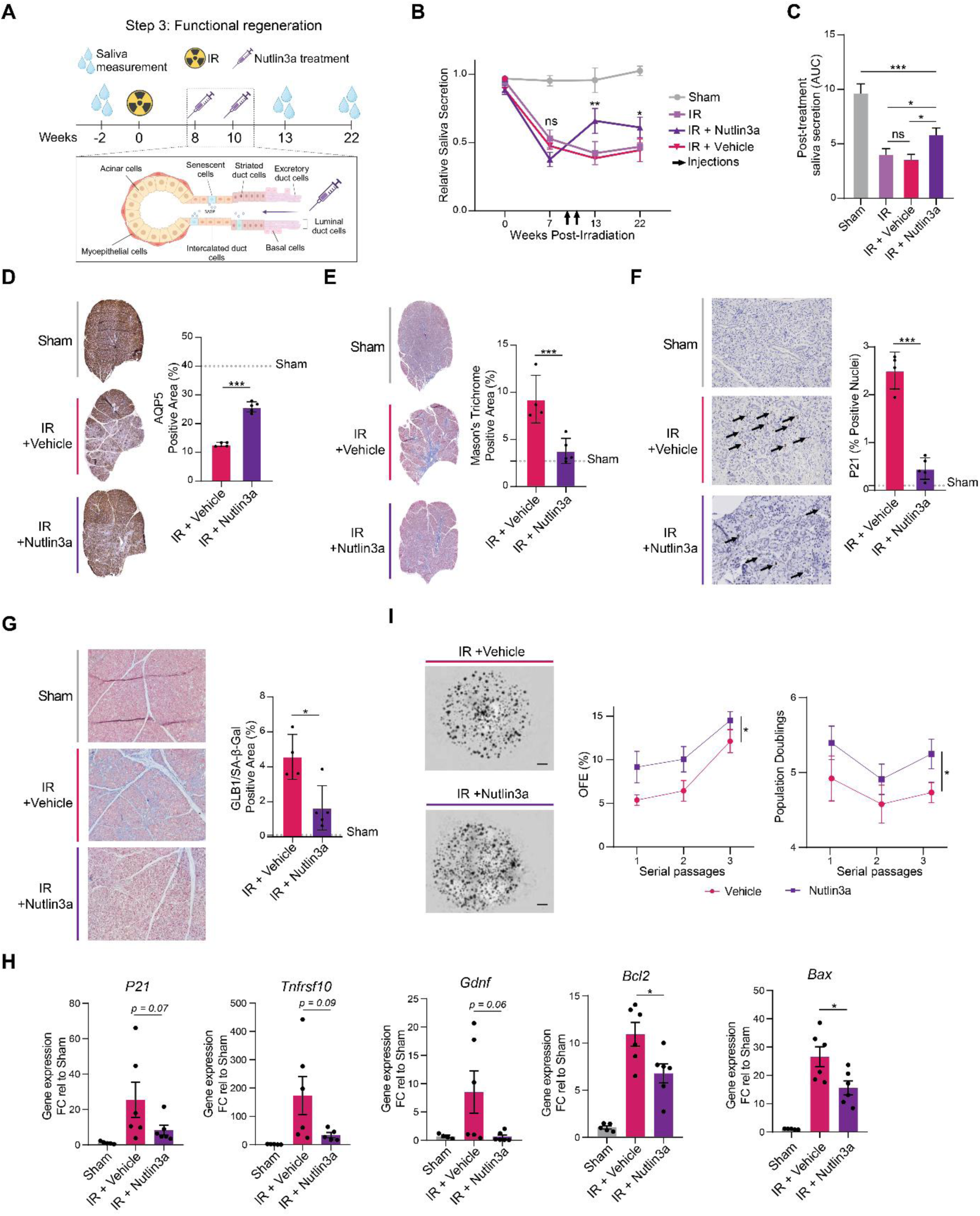
Local Nutlin3a treatment improves salivary gland function after radiation injury, reduces cell senescence and promotes stem cell function. (**A**) Experimental timeline showing 15 Gy irradiation to the neck area, followed by retroductal Nutlin3a administration at 8 and 10 weeks. (**B, C**) Saliva production was collected before irradiation (baseline) and after Nutlin3a treatment (n=6-8). A decrease in saliva production relative to baseline was seen at 7 weeks post-irradiation in all groups. Only Nutlin3a treatment produced a sustained partial recovery (IR + Nutlin3a), compared with non-injected mice (IR), and mice injected with vehicle alone (IR + Vehicle). (**D, E**) AQP5-positive acinar area and fibrosis at 13 weeks. (F, G) P21 positive cells and β-gal (GLB1) staining as senescence-associated readouts (**H**). Expression of senescence- and apoptosis-associated genes at 22 weeks post-irradiation. (**I**) Ex vivo organoid forming efficiency (OFE) of cells isolated from treated glands at the late endpoint, including serial passaging and population-doubling analyses. Data shown are mean ± sem, ns = not significant, \**p* < 0.05, \*\**p* < 0.01, \*\*\**p* < 0.001, \*\*\*\**p* < 0.0001, in unpaired *t*-tests.

Nutlin3a is generally well tolerated in mice, with previous studies employing oral twice daily doses of 200 mg/kg [32]. However, clinical trials using other MDM2 inhibitors have revealed significant gastrointestinal tract toxicity, including nausea, vomiting and diarrhea, often limiting the maximum tolerated dose [33]. Therefore, we decided to test if localized delivery would be a feasible strategy for obtaining a regional response and thereby minimize potential off-target and systemic side effects [34].

Salivary glands eject saliva into the mouth via excretory ducts, which if cannulated can be used for direct access via retroductal instillation/ injection [15]. As such, we delivered single doses of Nutlin3a (0.05 mg/kg) or vehicle alone, at 8- and 10-weeks post-irradiation and measured changes in saliva production at 3 weeks and 12 weeks after the last injection (or the equivalent to 13 weeks and 22 weeks post-irradiation).

Expectedly, saliva production was dramatically reduced at 7 weeks post-irradiation in all irradiated animal groups, before localized interventions. However, saliva production increased after retroductal injection in the Nutlin3a group (IR + Nutlin3a), but not in the vehicle treated group (IR + Vehicle), nor in the non-injected group (IR) at both 13 weeks and 22 weeks post-irradiation (**Fig. 5B,C**). Moreover, we observed an increased number of Aquaporin 5 (Aqp5)-positive acinar cells in the Nutlin3a treated group (**Fig. 5D**), accompanied with reduced fibrosis when compared to irradiated glands treated with vehicle at 13 weeks post-irradiation (**Fig. 5E**). These data indicate that localized treatment with Nutlin3a can improve the morphology and function of irradiated salivary glands as early as 2 weeks post-treatment.

Next, we sought to verify the effect of Nutlin3a on the presence of senescent cells *in vivo*. For this, we looked for P21^+^ cells, due to the pivotal role of P21 in cell senescence through its mediation of both acute and long-term cell cycle arrest in response to DNA damage. Accordingly, we observed a dramatic increase in P21^+^ cells at 13 weeks (**Fig. 5F**) and 22 weeks (Data not shown) after irradiation. Importantly, P21^+^ cells were virtually undetectable in sham-irradiated mice, indicating the persistence of P21^+^ cells several months after the single irradiation event. However, we observed a reduction in P21^+^ cells in the Nutlin3a treated group compared to vehicle treated mice at 13 weeks post-irradiation (**Fig. 5F)**, suggesting a clearance of P21^+^ cells after localized Nutlin3a treatment. Of interest, the decrease in P21^+^ cells in the Nutlin3a treated group was accompanied by a decrease in β-gal staining (Glb1) in irradiated glands suggesting a lower burden of senescent cells (**Fig. 5G**).

Next, we characterized the long-term effects of localized Nutlin3a on irradiated salivary glands, measured 12 weeks post-injection, or the equivalent of 22 weeks post-irradiation. At this timepoint saliva production continued to be partially restored in the Nutlin3a treated group. However, increased saliva production was accompanied by only modest histochemical improvements in Aqp5, fibrosis and markers of senescence, which failed to reach statistical significance at the 22-week endpoint (data not shown). To confirm the long-term impact of Nutlin3a in late-effects of radiation, we terminated all mice at this timepoint and measured cell senescence markers by RT-qPCR at this timepoint. Importantly, we observed lower Cdkn1a/P21, lower Tnfrsf10, and lower Gdnf in the Nutlin3a treated group (**Fig. 5H**). Similarly, we measured lower pro-apoptotic (Bax) and anti-apoptotic (Bcl2) signalling, suggesting lower apoptotic signalling. In addition, the long-term effects of Nutlin3a on stem cell capacity were tested *in vivo* by processing the extracted glands to isolate salivary gland cells and testing these for stem cell self-renewal capacity. As shown in **Fig. 5I**, we noted an increased organoid forming efficiency in irradiated salivary glands treated with Nutlin3a, relative to vehicle-treated irradiated glands. This improvement in organoid forming efficiency was retained after serial passaging, and was accompanied by an increase in population doubling, indicating higher proliferation. All together, these findings suggest an increase in stem cell capacity as late as 12 weeks after localized Nutlin3a treatment, suggesting its therapeutic potential as a novel strategy to ameliorate radiation-induced hyposalivation.

## DISCUSSION

This study establishes a two-step organoid screening strategy that evaluates candidate senolytics not only for preferential killing of radiation-damaged cells but also for their compatibility with tissue regeneration. Applying this approach to irradiated salivary gland organoids identified Nutlin3a as the candidate with the strongest combination of preferential apoptotic activity and recovery of organoid-forming capacity. Local administration of Nutlin3a subsequently improved salivary gland function *in vivo*, linking the *ex vivo* screening phenotype to a therapeutically relevant outcome.

The screening design addresses and important limitation of conventional senolytic discovery. Senescent-cell survival programs are heterogeneous and depend on cell type, senescence inducer, and tissue context [6, 27]. A compound that preferentially kills one senescent cell type may therefore be ineffective in another, while a drug that induces cell death may still compromise the regenerative compartment. It is thought that SASP factors released from senescent cells can reduce stemness and induce senescence of healthy cells in a paracrine manner, influencing the microenvironment by promoting the recruitment of inflammatory cells and via the induction of fibrosis and tissue remodelling. In turn, both senescent cells, and infiltrated immune cells may exacerbate local chronic inflammation. Given the pro-inflammatory roles of senescent cells, we reasoned that apoptotic cell death, generally recognized as anti-inflammatory and immunosuppressive [35–37], may improve the resolution of local tissue inflammation. This may be accomplished not only by removing the inflammatory senescent cells present in the tissue and reducing their secretion of chemoattractant molecules, but also by having an immunosuppressive effect on immune cells that phagocytose these apoptotic cells [37].

In our RNA-seq experiments comparing irradiated SG organoids with controls, we observed the upregulation of both pro-apoptotic and anti-apoptotic members of the Bcl-2 family of proteins. The highest overexpressed genes were Bmf (Bcl-2 Modifying Factor) and Bik (Bcl-2 Interacting Killer), two proteins that can bind Bcl-2 and function as apoptotic activators. Interestingly, induction of cell death by these proteins is mediated by an entirely Bax-dependent mitochondrial pathway [19]. Bax in turn, was also identified as an upregulated gene in our RNA-seq experiment, and its induction was validated by RT-qPCR in irradiated organoids and further upregulated upon Nutlin3a treatment. This propensity to engage the mitochondrial apoptotic pathway following irradiation may explain why irradiated organoids were more susceptible to inhibition of Bcl-2, Bcl-Xl, and Bcl-w (all of which were upregulated upon irradiation), by the small molecule inhibitor ABT-263 [24].

The priming to apoptosis we observed in irradiated organoids, may explain why irradiated organoids are more susceptible to MMD2 inhibition by Nutlin3a. Indeed, a recent study using an unbiased gene knockdown approach in (senescent) embryonic fibroblasts proposed the active suppression of P53 in senescent cells to remain viable, including its modulation by MDM2. Interestingly, despite an emphasis on the transcriptional upregulation of pro-apoptotic target genes upon MDM2 inhibition, its effect on anti-apoptotic proteins was not explored. However, we believe the downregulation of the anti-apoptotic proteins Bcl-2 and Birc5/Survivin upon MDM2 inhibition also contributes to the effectivity of Nutlin3a-induced apoptosis in senescent cells.

The observed downregulation of Bcl-2 and Birc5/Survivin is likely due to p53 activation by Nutlin3a, as both genes are transcriptionally repressed by wild-type p53 and have p53-dependent negative response elements in their promoter regions [38, 39]. Accordingly, downregulation of Bcl-2 and Survivin was observed in both non-irradiated (sham) and irradiated cells treated with Nutlin3a. However, cell death was mainly observed in irradiated organoids, presumably due to a bigger dependence of irradiated cells on these anti-apoptotic proteins to restrain their exacerbated pro-apoptotic signaling. Therefore, the downregulation of anti-apoptotic proteins such as Survivin by Nutlin3a may synergize with upregulated pro-apoptotic (such as Bax) and result in increased cell death of radiation-induced senescent cells.

In addition, treatment with Nutlin3a resulted in increased organoid forming efficiency of irradiated organoids, both *ex vivo* and *in vivo,* suggesting a partial rescue of the deleterious effects of irradiation on stem cell expansion. Importantly, enhanced stem cell expansion was accompanied by a functional rescue in saliva production that persisted to 22 weeks. It remains to be tested whether these effects persist beyond this timepoint, or whether additional rounds of Nutlin3a treatment, may re-prompt a larger regenerative response.

In summary, integrating selective cell-death measurements with a functional regenerative readout provides a practical strategy for identifying tissue-adapted senolytic interventions. In irradiated SG organoids this approach identified transient MDM2 inhibition as a means of exploiting an apoptosis-primed state, and local Nutlin3a treatment translated into sustained improvement of salivary gland function in vivo. More broadly, organoid screes that incorporate both senolytic activity and regenerative competence may help prioritize senotherapeutics for radiation injury and other tissue-specific settings.

## ACKNOWLEDGEMENTS

This project was funded by KWF project 12458.

## Notes

### Competing Interest Statement

The authors have declared no competing interest.

