## Supplemental Tables and Figures for "A two-step organoid screen identifies transient p53 activation as a senolytic strategy for radiation-induced salivary gland dysfunction"

651 **Supplementary Table 1 – Drug concentrations used**

| *Name* | *Concentration* |  |  |  |  |
| --- | --- | --- | --- | --- | --- |
|  | [1] | [2] | [3] | [4] | Unit |
| *ARV825* | 50 | 100* | 200 | 400 | nM |
| *17DMAG* | 50 | 100 | 200 | 400 | nM |
| *Geldanamycin* | 50 | 100 | 200 | 400 | nM |
| *Dasatinib + Quercetin (D+Q)* | 1+5* | 5+25 | 10+50 | 20+100 | nM+μM |
| *TH588* | 10* | 20 | 40 | 80 | μM |
| *2-DG* | 5 | 10 | 20 | 40 | mM |
| *Fisetin* | 2,5 | 5 | 10 | 20 | μM |
| *TRAIL DHER* | 125 | 250 | 500 | 1000 | ng/ml |
| *Nutlin3a* | 2 | 4 | 8* | 16 | μM |
| *ABT-263* | 30 | 75 | 150 | 300* | nM |
| *Staurosporine (STS)* | 62.5 | 125 | 250 | 500 | nM |

652 The concentrations used for self-renewal assays of Figure 2 are marked with an asterisk.

653

654 **Supplementary Table 2 – List of primers used for gene expression analysis**

| *Gene* | *Forward primer (5' -> 3')* | *Reverse primer (5' -> 3')* |
| --- | --- | --- |
| *Cdkn1a/P16* | GAACTCTTTCGGTCGTACCC | CGAATCTGCACCGTAGTTGA |
| *Tnfrsf10* | GCTGTGTCTGTGGCTGTGACTT | TCCTCATCCGTCTTTGAGAAGC |
| *Gdnf* | CGCTGACCAGTGACTCCAATATG<br>C | TGCCGCTTGTTTATCTGGTGACC |
| *Cxcl10* | CCAAGTGCTGCCGTCATTTTC | GGCTCGCAGGGATGATTTCAA |
| *P53inp1* | GAGACTCGAGCTTGGTTCACCGCTCT<br>CACT | GAGAGGATCCTCGTCTGGTAGCTCTT<br>CGGT |
| *Cdkn1a/P21* | AGGCAGACCAGCCTGACAGAT | TCCTGACCCACAGCAGAAGAG |
| *Lif* | TGAAAACGGCCTGCATCTA | GACCTTCATTATGGGCTGGA |
| *Mdm2* | CCAACCATCGACTTCCAGCAGCATT | GATTGGCTGTCTGCACACTGGG |
| *Puma* | TGGAGGGTCATGTACAATCTCTT | GTTGGGCTCCATTTCTGG |
| *Bim* | GGAGACGAGTTCAACGAAACTT | AACAGTTGTAAGATAACCATTTG<br>AGG |
| *Bak* | CCAGCTGATGCCACTCTTAAA | GGAATGCCTACGAACTCTTCA |
| *Bax* | GTGAGCGGCTGCTTGTCT | GTGGGGGTCCCGAAGTAG |
| *Bcl-2* | AGTACCTGAACCGGCATCTG | GGGGCCATATAGTTCCACAAA |
| *Birc5/Survivin* | CCCGATGACAACCCGATA | CATCTGCTTCTTGACAGTGAGG |
| *Bcl-XL* | AGTGCAGGATTGGATGGTG | CCCGTATAGAGCTGTGAACTCC |
| *Mcl-1* | GGTATTTAAGCTAGGGTCATTTGAA | TGCAGCCCTGACTAAAGGTC |
| *Ywhaz* | TTACTTGGCCGAGGTTGCT | TGCTGTGACTGGTCCACAAT |

655

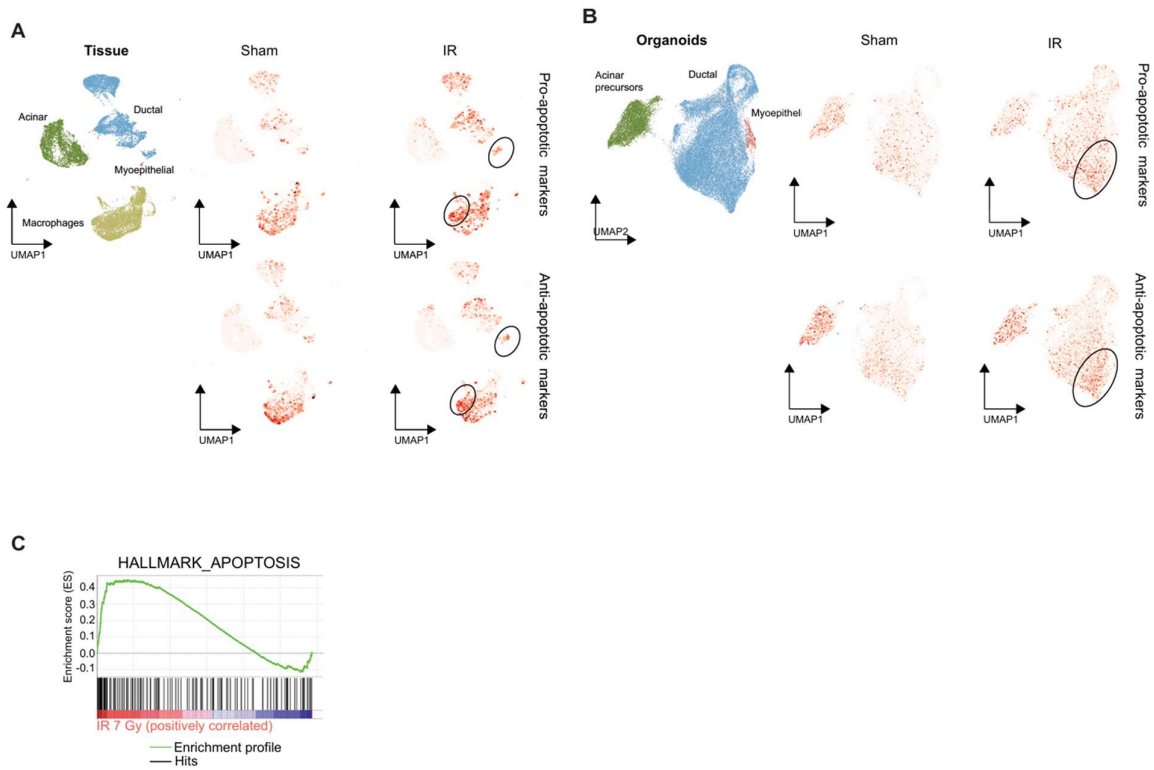

**Supplementary Figure 1. Increased pro- and anti-apoptotic signaling in irradiated salivary gland tissue and organoids.** Average expression of both pro-apoptotic (Bax, Bak1, Bcl2l11, Bbc3) and anti-apoptotic (Bcl2, Bcl2l1, Mcl1, Mdm2) markers after irradiation in suspected senescent populations of salivary gland tissue (A, 10 Gy, [22]) and organoids (B, 7 Gy, [16]). Gene set enrichment analysis of bulk transcriptomic data from (sham)-irradiated salivary gland organoids confirms a positive correlation of the apoptosis hallmark in (7 Gy) irradiated organoids.

664

**A**

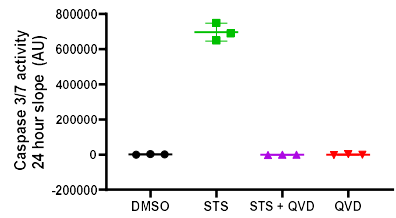

**B**

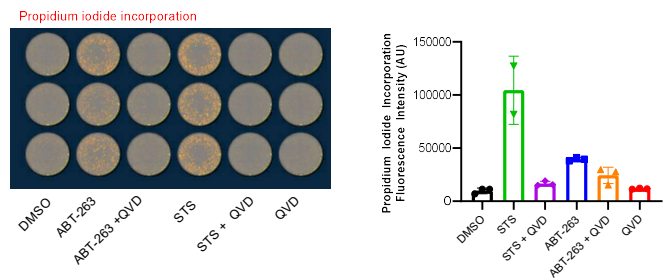

665

666 **Supplementary Figure 2. Evaluating drug cytotoxicity in salivary gland organoids.** (A) Increased cytotoxicity of STS compared
667 to DMSO was measured by CellEvent™ Caspase-3/7 Green dye in salivary gland organoids. The plot shows the change in
668 fluorescence intensity from Caspase-3/7 activity after 24 hours of treatment with staurosporine (STS, 0.5 μM) in the presence
669 or absence of the pan-caspase inhibitor QVD-Oph (QVD, 10 μM), which inhibited a fluorogenic response. B) Propidium iodide
670 incorporation was used as an additional readout for cell death. The fluorescence intensity of organoids treated with
671 staurosporine or ABT-263 (0.3 μM) was calculated. The presence of QVD inhibited fluorogenic responses in drug treated
672 organoids, suggesting caspase dependent cell death. Graphs show mean fluorescence intensity ± SD.

673

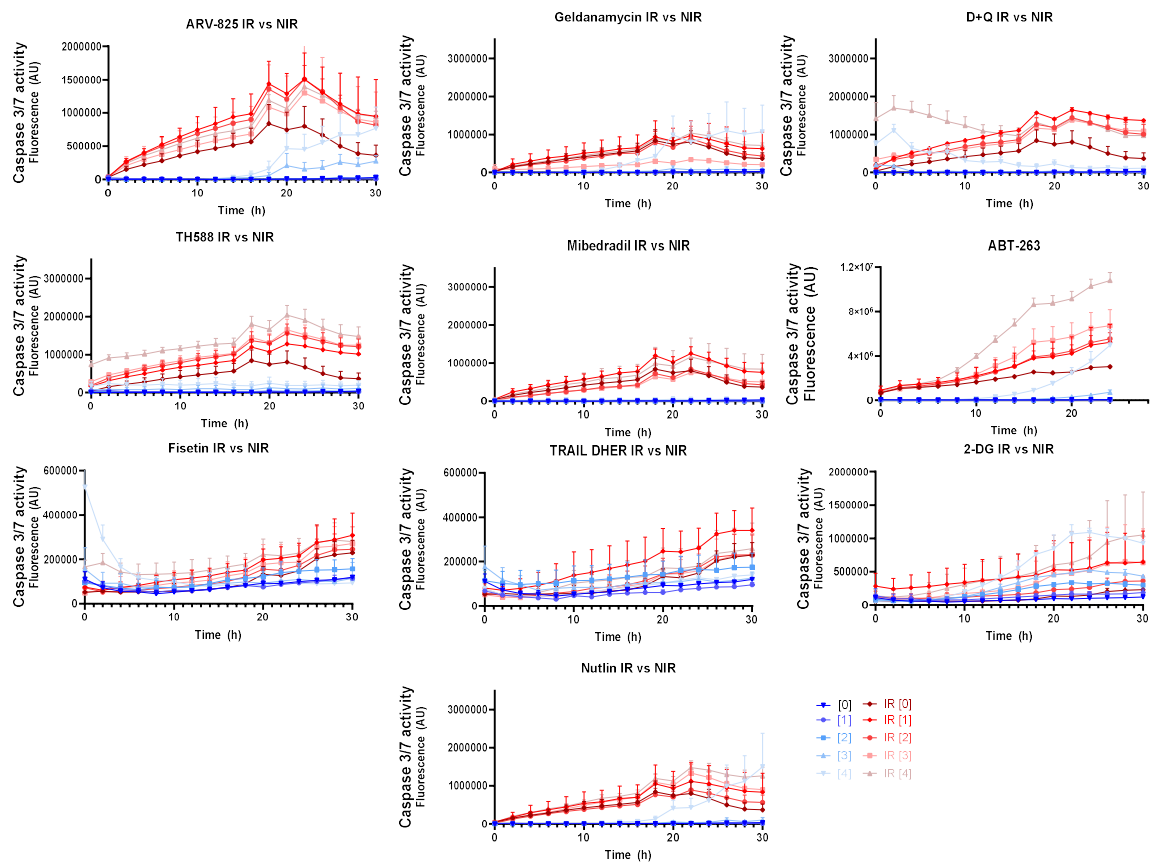

674

675 **Supplementary Figure 3. Time resolved assays illustrating Caspase-3/7 activity after treatment of different**
676 **drugs in (non-)irradiated organoids.** Graphs show mean fluorescence intensity ± SD from representative
677 experiments performed in triplicates. The numbers between brackets refer to the different concentrations used
678 as indicated in Supplementary Table 1.
